# Genome assembly, annotation and comparative analysis of the cattail *Typha latifolia*

**DOI:** 10.1101/2021.08.23.457420

**Authors:** Shane D. Widanagama, Joanna R. Freeland, Xinwei Xu, Aaron B.A. Shafer

## Abstract

Cattails (*Typha* species) comprise a genus of emergent wetland plants with a global distribution. *Typha latifolia and T. angustifolia* are two of the most widespread species, and in areas of sympatry can interbreed to produce the hybrid *Typha* x *glauca*. In some regions the relatively high fitness of *T*. x glauca allows it to outcompete and displace both parent species, while simultaneously reducing plant and invertebrate biodiversity, and modifying nutrient and water cycling. We generated a high-quality whole genome assembly of *T. latifolia* using PacBio long-read and high coverage Illumina sequences that will facilitate evolutionary and ecological studies in this hybrid zone. Genome size was 287 Mb and consisted of 1,189 scaffolds, with an N50 of 8.706 Mb; 43.84% of the genome were identified as repetitive elements. The assembly has a BUSCO score of 96.03%, and 27,432 genes and 2,700 RNA sequences were putatively identified. Comparative analysis detected over 9,000 shared orthologues with related taxa and phylogenomic analysis supporting *Typha latifolia* as a divergent lineage within Poales. This high-quality scaffold-level reference genome will provide a useful resource for future population genomic analyses and improve our understanding of *Typha* hybrid dynamics.

## INTRODUCTION

Cattails (*Typha* spp.) are aquatic macrophytes that are essential components of wetlands around the world (reviewed in Bansal et al., 2019). These macrophytes often dominate ecosystems through a combination of rapid growth, large size, and sexual and asexual reproduction (Yeo, 1964; Andrews and Pratt, 1978; Miller and Fujii, 2010). Cattails are vital to many wetlands where they cycle nutrients, provide habitat, and aid in bioremediation (Grosshans, 2014; Svedarsky et al., 2016; Bonanno and Cirelli, 2017). However, cattails can also dominate wetlands and in recent decades invasive cattails have been identified in numerous regions, often following anthropogenic changes that have altered water cycles and increased nutrient loads in wetlands (reviewed in Zedler and Kercher, 2004; Bansal et al., 2019). Invasive cattails often form monotypic stands with deleterious effects on local plants and animals, wetland water cycling, and biogeochemical cycles (Yeo, 1964; Grace and Harrison, 1986; Farrer and Goldberg, 2009; Lawrence et al., 2016).

*Typha latifolia* is likely the most widespread cattail species, occurring on every continent except Antarctica (Smith, 1987). *Typha angustifolia* is also widespread throughout the temperate northern hemisphere (Grace and Harrison, 1986), including North America where it was likely introduced from Europe several centuries ago (Ciotir et al., 2013, 2017). In some regions of sympatry *T. latifolia* and *T. angustifolia* interbreed to produce the hybrid *Typha* x *glauca* (Grace & Harrison, 1986; Ciotir et al., 2017; Bansal et al., 2019). In regions surrounding the Laurentian Great Lakes and St. Lawrence Seaway in North America, *T*. x *glauca* exhibits heterosis (Bunbury-Blanchette et al., 2015; Zapfe and Freeland, 2015), and is more abundant than its parental species (Kirk et al., 2011; Freeland et al., 2013; Pieper et al., 2020). In addition to displacing both parental species, invasive *T*. x *glauca* reduces native plant and invertebrate biodiversity (Lawrence et al., 2016; Tuchman et al., 2009), and alters nutrient cycling and community structure in wetlands (Tuchman et al., 2009; Larkin et al., 2012; Geddes et al., 2014; Lishawa et al., 2014; Lawrence et al., 2017).

Although dominant in some regions of North America, *T*. x *glauca* is uncommon in other regions where the parental species are sympatric, including Europe (Ciotir et al., 2017), eastern Canada (Freeland et al., 2013), and China (Zhou et al., 2016). It is not well understood why hybrids are dominant in some regions but not others, although Tisshaw et al., (2020) suggested that hybrids may be limited in coastal wetlands because they have difficulty germinating in salt-rich environments. In addition, although advanced-generation and back-crossed hybrids have been experimentally generated (Pieper et al., 2017) and identified in natural populations (e.g. Kirk et al., 2011; Freeland et al., 2013; Pieper et al., 2020), it has not been possible to differentiate advanced-generation and backcrossed hybrids based on the small number of species-specific molecular markers that are currently available (Kirk et al., 2011; Snow et al., 2010). Morphology-based assessments are also unreliable due to overlapping phenotypes (Tangen et al., 2021).

A genome-wide suite of SNPs specific to one or the other parent species would greatly facilitate investigations into the dynamics of the *T*. x *glauca* hybrid zone, which is now expanding westwards from the Great Lakes region into the Prairie Pothole Region of Canada and the USA (Tangen et al., 2021). Genome characterization would also facilitate investigations into local adaptation, introgression, and hybrid dynamics; this in turn may inform future management of *T. latifolia*, which is predicted to experience a dramatically reduced distribution following climate change (Xu et al., 2013), likely to the detriment of wetlands throughout its widespread native distribution. Here we report the genome assembly, annotation and analysis of *T. latifolia*, the first fully sequenced species in the family Typhaceae.

## MATERIALS & METHODS

### Sampling and Sequencing

Leaves of a known *T. latifolia* plant (Figure 1) were taken from an individual grown at Trent University’s (Ontario, Canada) greenhouse and ground in liquid nitrogen. DNA was immediately extracted using an E.Z.N.A Plant DNA Kit (Omega Bio-Tek, Inc. Georgia USA) following the manufacturer’s instructions for frozen material and eluted in a final volume of 100 μl. DNA quality was assessed on a Tapestation (Agilent Technologies, California USA) before being sent to The Centre for Applied Genomics, Toronto, Ontario, for sequencing. Paired-end reads were sequenced on a single lane of an Illumina HiSeqX system (Illumina, Inc. California USA). Long-reads (LR) were sequenced on one SMRT HiFi cell using a PacBio Sequel II system (Pacific Biosciences of California, Inc. California USA).

**Figure 1.**
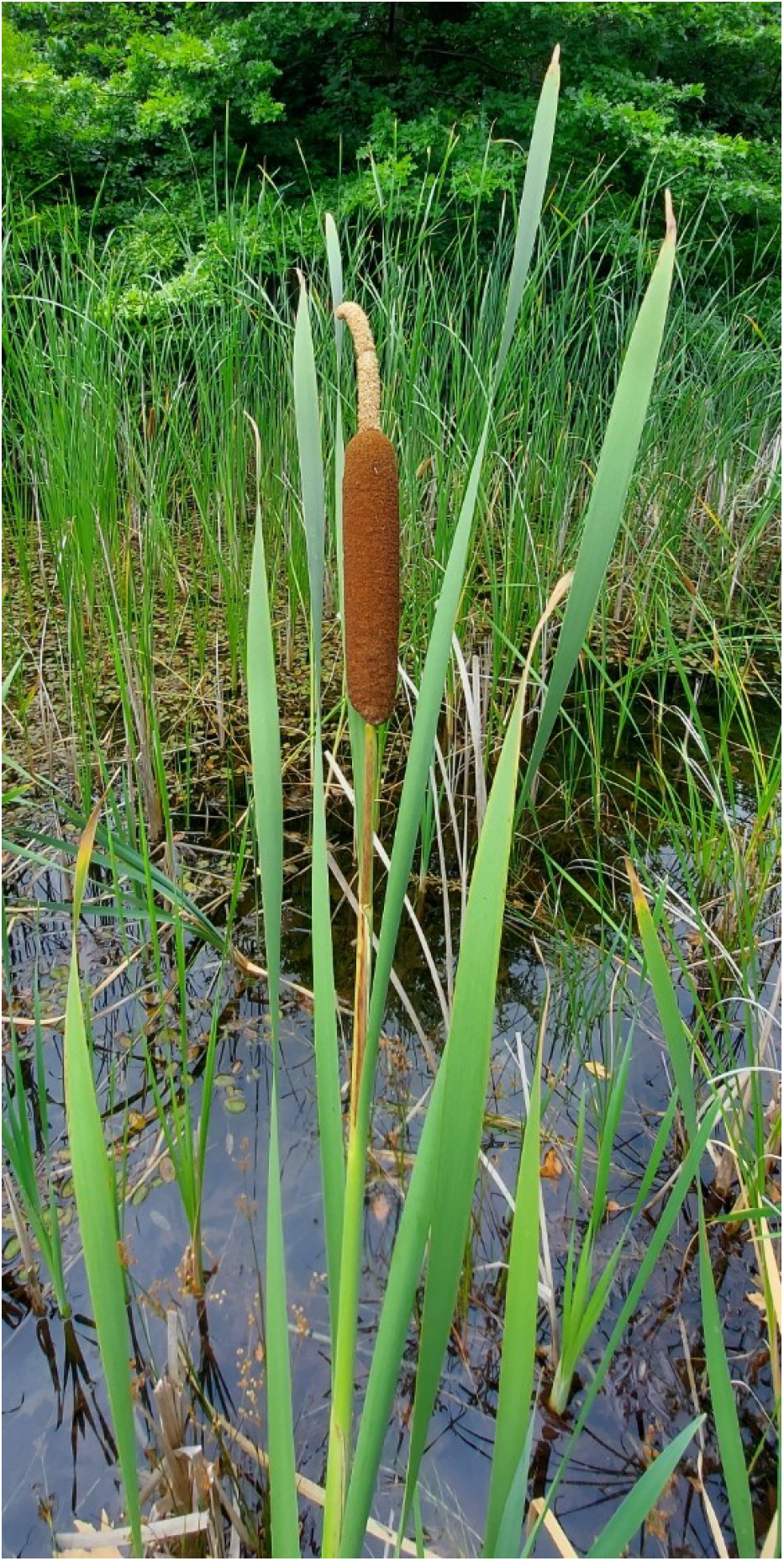
Broadleaf cattail (*Typha latifolia*). Photo by Joanna Freeland.

### Genome Assembly

Adapters and low-quality bases were trimmed from paired-end short reads (SR) using Trimmomatic −v0.36 (Bolger et al., 2014). The optimal k-mer and genome size from the short reads was estimated using Kmergenie −v1.7051 (Chikhi & Medvedev, 2014). We then applied a four-step hybrid assembly: 1) SR assemblies were generated using multiple pipelines: ABySS −v2.2.4 (Jackman et al., 2017) with default settings, 32 threads, a 125 GB total memory; SOAPdenovo2 −r240 (Luo et al., 2012) using default settings, 16 threads, 400 GB of total memory; and Platanus −v1.2.4 (Kajitani et al., 2014) using default settings, 32 threads, a 343 GB memory limit, a total memory of 375 GB. We also downsampled the SR data to 20% because really high coverage can be problematic for de Bruijn graphs (Richards & Murali, 2015). The SR assembly with highest quality and completeness was selected by calculating contig N50, total contig length as a percentage of estimated genome size, and the number and percentage of contigs longer than 50 Kb. 2) SR contigs were then aligned to raw long-reads (LR) to create scaffolds using DBG2OLC −v20160205 (options: k 17, AdaptiveTh 0.001, KmerCovTh 2, MinOverlap 20, RemoveChimera 1) (Ye et al., 2016). 3) LR contigs were assembled using Canu −v2.0 (Koren et al., 2017) using the expected genome size inferred from Kmergenie, a maximum memory of 124 GB, and a maximum of 32 threads. And 4) the hybrid assembly (step 2) was merged with Canu LR contigs (step 3) using QuickMerge −v0.3.0 (Chakraborty et al., 2016) following the 2-step strategy described by Solares et al., (2018). Pilon −v1.23.0 (Walker et al., 2014) was used to polish the genome by mapping the Illumina SR back to the genome, thereby correcting base errors and small misassemblies.

### Genome Evaluation and Annotation

The *T. latifolia* hybrid genome was assessed for completeness using Benchmarking Universal Single-copy Orthologs (BUSCO) – v3.0.2 (Simão et al., 2015). BUSCOs of the lineage Liliopsida were assessed from OrthoDB release 10 (Waterhouse et al., 2013). We aligned available chloroplast and mitochondrial genomes (Table S1) to our assembly using NUCmer −v3.23 (Kurtz et al., 2004) to identify plastid genomes; scaffolds with long matches (>5000 bp) were extracted and further validated on NCBI Blast. Structural and functional annotation was done in the GenSAS −v6.0 annotation pipeline (Humann et al., 2019). We masked interspersed and simple repetitive elements throughout the genome using a database developed through repeat modeler −v2.0.1 (Smit & Hubley, 2015) in conjunction with repeat masker −v4.0.7 with the NCBI/rmblast search engine, and quick sensitivity (Smit, Hubley & Green, 2015). The hardmasked version of the genome sequence was used for feature prediction. Available RNA-seq data from *T. angustifolia* (SRR15541138) were mapped to the genome using hisat2 −v2.1.0 (Kim et al., 2019) and gene prediction with the RNA-seq alignments was performed by the BRAKER2 −v2.1.1 pipeline, which uses Augustus and GeneMark-ET (Lomsadze et al., 2014, Brůna et al., 2021). ncRNA was predicted using tRNAscan-SE −v2.0 (Lowe & Eddy 1997) and Infernal −v1.1.3 (Nawrocki & Eddy 2013). Refinement of the official gene set was performed with an available *T. latifolia* transcriptome (Moscou, 2017) using PASA −v2.3.3 (Haas et al., 2003).

### Comparative Genomics

Single-copy orthologous genes from five other species were identified using OrthoVenn2 with an e-value cutoff of 1e-5, an inflation value of 1.5, and the annotation, protein similarity network, and cluster relationship network enabled (Xu et al., 2019). OrthoVenn2 performs all-against-all genome-wide protein comparisons and groups genes into clusters with the Markov Clustering Algorithm, where a cluster is made up of orthologs and paralogs (Wang et al., 2015). Here we selected *Oryza sativa Japonica, Brachypodium distachyon*, and *Sorghum bicolor* from the family Poaceae, and *Ananas comosus* from the family Bromeliaceae*. Arabidopsis thaliana* from the Brassicaceae family was chosen as the dicot outgroup. All proteomes were from the Ensembl database release 104 (Howe et al., 2021). The identified orthologous sequences were then aligned using MAFFT −v7.741 (Katoh & Standley, 2013) and concatenated into single sequences by species using SeqKit −v0.15.0 (Shen et al., 2016). Maximum likelihood phylogenetic analysis was performed using RAxML - v8.2.12 (Stamatakis, 2014) with the PROTGAMMAAUTO substitution model (Darriba et al., 2012). The phylogenetic tree divergence times were estimated using MCMCTree from the PAML package −v4.9j (Yang 1997, 2007), and was calibrated using the divergence time between *Sorghum* and *Oryza* (42-52 Mya) (Kumar et al., 2017).

## RESULTS

### Sequencing data and genome assembly

A total of 138.6 Gb of raw 151 bp Illumina reads were sequenced (Table S2). Quality filtering and trimming removed 6.5 Gb of sequence data. A recommended k-mer size of 101 bp and estimated genome size of 257 Mb was estimated from the short-read data (Figure S1). Long-read data from the Pacbio Sequel II generated 86.8 Gb of raw data: this included 7,244,218 subreads with a mean length of 11,978.2 bp. All sequencing reads were deposited in the NCBI Sequence Read Archive (Accession No PRJNA751759). The ABySS assembly that used 100% of Illumina reads had the most contiguous genome and was used for the assembly (Table S3). This AbySS SR assembly had a N50 of 0.011 Mb, 365,565 contigs, and 362 contigs longer than 50 Kb (Table S3). The DBG2OLC assemblies (ABySS contigs + raw long reads) improved the assembly statistics but resulted in a smaller than expected genome (Table 1); here we calculated N50 of 0.132 Mb, 1,840 contigs, and 1,445 contigs longer than 50 Kb (Table S4). The LR Canu assembly had an N50 of 8.706 Mb, and contained 1,189 contigs, 821 of which were longer than 50 Kb (95.54%) (Table 1). The final merged and polished hybrid genome assembly – DGB2OLC assembly 1 + Canu - was 287 MB, with an N50 of 8.706 Mb, and consisted of 1,189 scaffolds (Table 1). The GC content was 38.11% (Table 1). The polished genome has been deposited in the NCBI genome database.

**Table 1.** Genome assembly statistics of our four-step hybrid assembly. Short read (SR) contig assembly consists of contigs assembled from 100% of the Illumina reads using the ABySS assembler. The DBG2OLC assembler combined the 100% ABySS contigs and PacBio long reads. The long read (LR) contig assembly was generated from only PacBio long reads using Canu. The merged polished scaffolds are from merging the DBG2OLC assembly and LR contigs and were polished using Pilon.

|  | **SR contig**<br>**assembly –**<br>**step 1** | **DBG2OLC**<br>**contig**<br>**assembly –**<br>**step 2** | **LR contig**<br>**assembly –**<br>**step 3** | **Merged**<br>**polished**<br>**scaffolds –**<br>**step 4** |
| --- | --- | --- | --- | --- |
| **Genome Size**<br>**(Mb)** | 263.74 | 193.16 | 287.63 | 287.80 |
| **Contigs/**<br>**Scaffolds** | 365,565 | 1,840 | 1,190 | 1,189 |
| **N50 / L50** | 11.43 Kb /<br>5,314 | 132.07 Kb /<br>412 | 8.71 Mb / 13 | 8.71 Mb / 13 |
| **N90 / L50** | 201 bp /<br>193,744 | 52.95 Kb /<br>1,358 | 58.92 Kb / 530 | 58.94 Kb / 529 |
| **Max sequence**<br>**Length** | 154.73 Kb | 934.40 Kb | 18.70 Mb | 18.69 Mb |
| **Scaffolds > 10**<br>**Kb** | 6,048 | 1,833 | 1,140 | 1,139 |
| **Scaffolds > 25**<br>**Kb** | 1,759 | 1,785 | 1,127 | 1,126 |
| **Scaffolds > 50**<br>**Kb** | 362 | 1,445 | 821 | 821 |
| **% of scaffolds**<br>**> 50 Kb** | 9.46 | 92.34 | 95.54 | 95.55 |
| **GC content**<br>**(%)** | 38.40 | 38.50 | 38.11 | 38.11 |

### Assembly quality and Annotation

Approximately 96.03% of BUSCOs were identified (3,148 of 3,278) in the assembly. Of these, 3,025 BUSCOs were complete (92.28%), and 123 were fragmented. Of the complete BUSCOs, 2,461 were single-copies, and 564 had duplicates. Repeats represented 43.84% of the genome (see breakdown in Figure 2). The chloroplast genome was detected and split on two scaffolds, while the entire mitochondrial genome was assembled (Table S5). Total repeat content of the genome is consistent with other Poales genomes (Kawahara et al., 2013, Ming et al., 2015, Redwan et al., 2016): long interspersed nuclear elements (LINEs) comprised 1.22% of the genome, while short interspersed nuclear elements (SINEs) were not detected in the genome (Table 2). The annotation pipeline produced 27,432 genes, which coded for 34,911 proteins and 34,974 mRNA sequences. 2,095 rRNA, 502 tRNA, and 214 miRNA sequences were putatively identified, which are similar to sequence counts in related plants (Table S6). No snRNA were identified.

**Figure 2.**
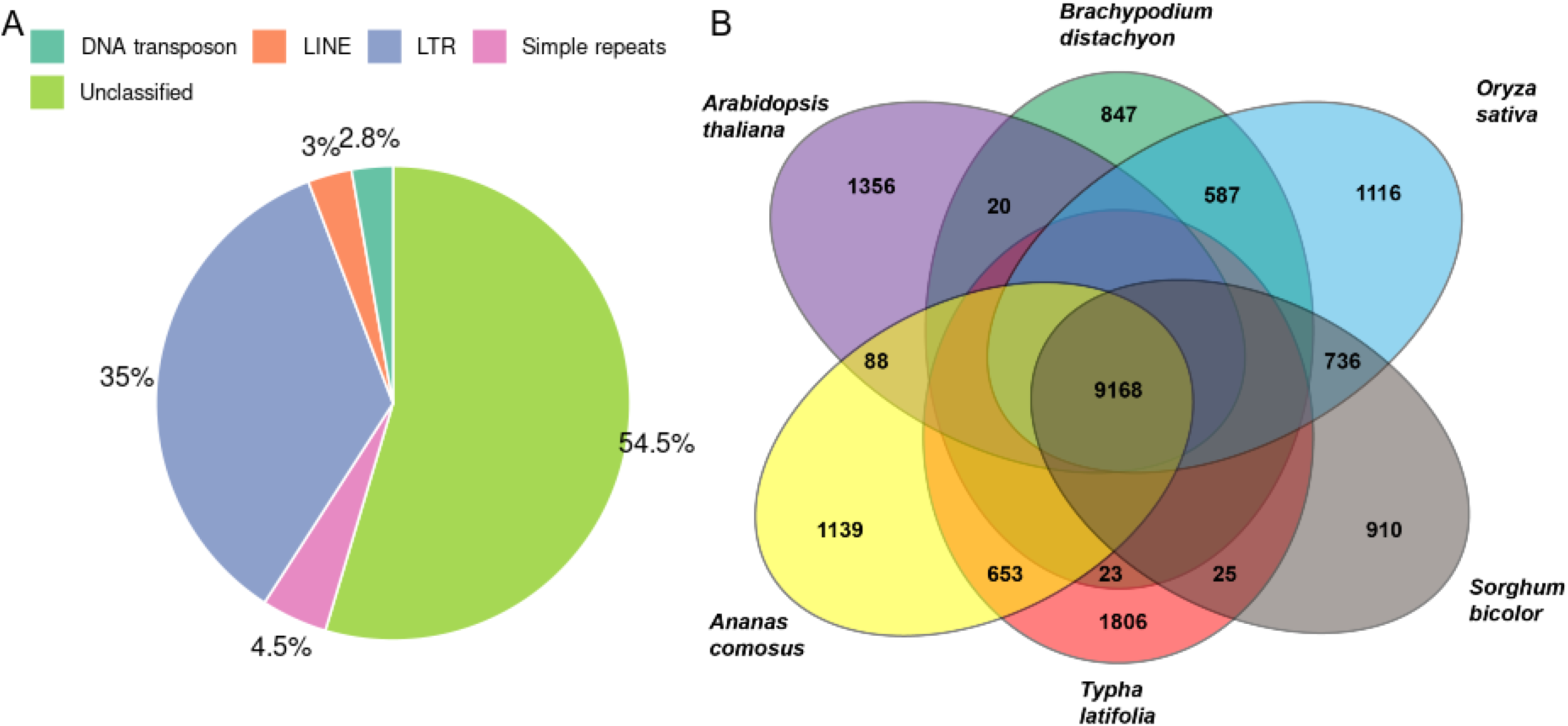
a) Percentages of repeat types masked in the *T. latifolia* genome. Types of repeats include DNA transposons, long interspersed nuclear elements (LINEs), long terminal repeats (LTRs), unclassified repeats, and simple repeats. b) Venn diagram of orthologous gene clusters among the broadleaf cattail (*Typha latifolia*), pineapple (*Ananas comosus*), thale cress (*Arabidopsis thaliana*), stiff brome (*Brachypodium distachyon*), rice (*Oryza sativa Japonica*), and broom-corn (*Sorghum bicolor*). Only the numbers of ortholog clusters of adjacent species, those common to all species, and those unique to each species are labelled.

**Table 2.** Summary of repeats masked in the *Typha latifolia* genome.

|  | Length (bp) | Percentage of genome (%) |
| --- | --- | --- |
| SINE | 0 | 0 |
| LINE | 3,499,426 | 1.22 |
| LTR elements | 44,172,983 | 15.35 |
| DNA elements | 3,735,882 | 1.30 |
| Unclassified | 68,736,281 | 23.88 |
| Small RNA | 0 | 0 |
| Satellites | 0 | 0 |
| Simple repeats | 5,689,134 | 1.98 |
| Low complexity | 670,871 | 0.23 |
| Total | 126,178,695 | 43.84 |

### Comparative Genomic Analyses

Ortholog clustering analysis revealed 9,168 gene families were shared among *T. latifolia, A. comosus, A. thaliana, B. distachyon, O. sativa, and S. bicolor* (Figure 2). 1,806 gene families were found to be unique to *Typha latifolia* (Figure 2). We aligned 1,900 single-copy gene clusters to conduct phylogenomic analysis (Figure 3). The phylogenomic tree generated supports a divergent *Typha* position in Poales (Figure 3), with *Typha* forming a separate clade with pineapples (*Ananas comosus*). We estimated the bromeliad lineage (*Typha-Ananas*) to be approximately 70 million years old.

**Figure 3.**
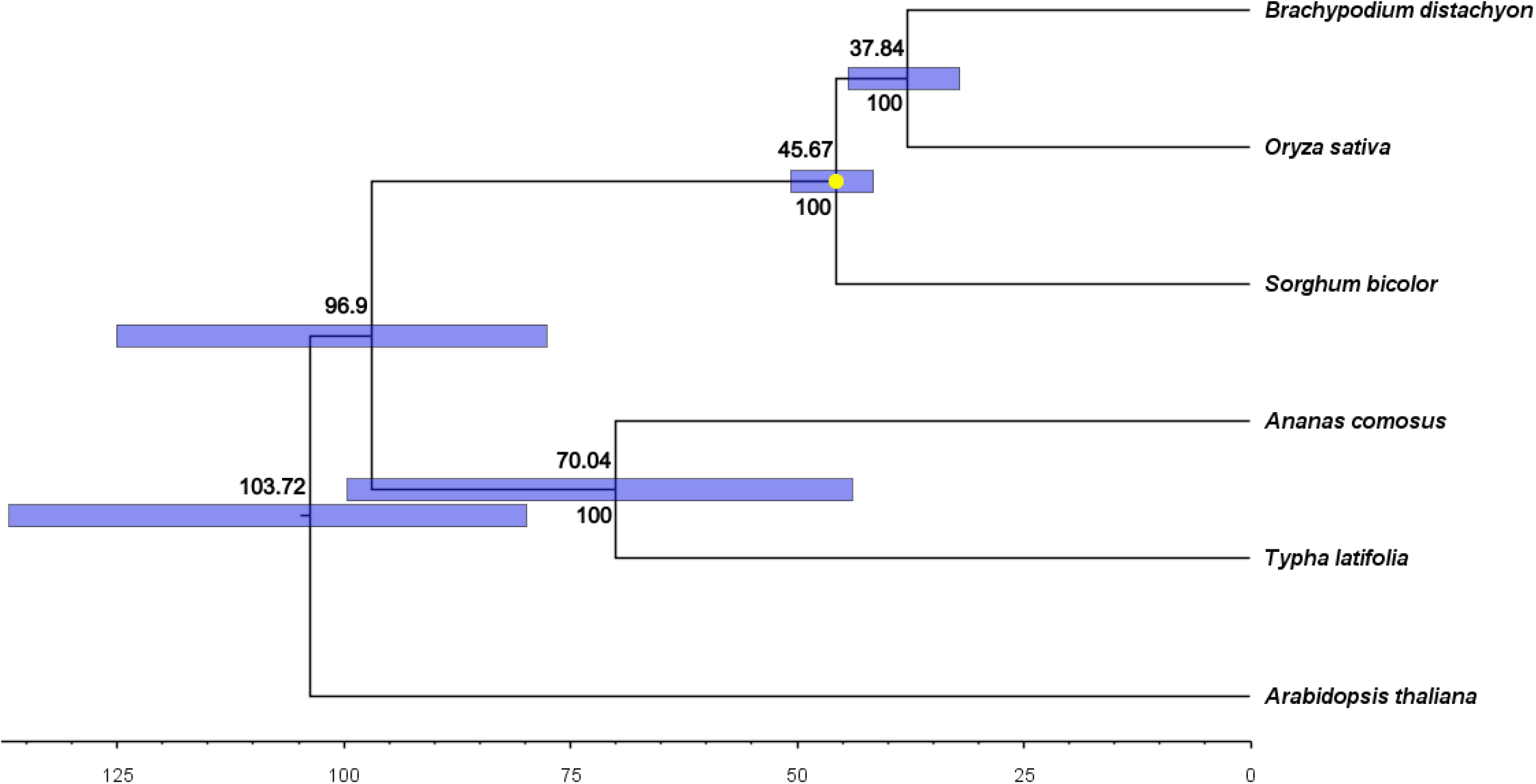
Phylogenetic tree with divergence times, based on the alignment of 1,900 single-copy gene clusters. 95% credible divergence times are shown as blue bars and were estimated using MCMCTree. Divergence times and bootstrap values are shown above and below the nodes, respectively. The yellow dot indicates the calibration point.

## DISCUSSION

The Typhaceae family (order Poales) is diverse, comprising over 50 recognized species (Christenhusz & Byng, 2016). Species in this family are essential components of marshes and wetlands around the world (reviewed in Bansal et al., 2019), and play a role in both bioremediation and the production of biofuel material (Grosshans, 2014; Svedarsky et al., 2016; Bonanno and Cirelli, 2017). This is the first whole genome sequence assembled from a member of the Typhaceae family and adds to the previously characterized chloroplast genome sequences of *Typha latifolia, T. orientalis*, and *Sparganium stoloniferum* (Guisinger et al., 2010; Su et al., 2019; Liu et al., 2020). The assembly and annotation of the Cattail genome, which we note was a key recommendation from a diverse team of *Typha* researchers (Bansal et al., 2019), will be an important tool in wetland management.

We explored a variety of short read assemblers given the varying quality of outputs seen in Assemblathon metrics (Bradnam et al., 2013). Of the three SR assemblers used, ABySS produced longer, fewer contigs and scaffolds, which is consistent with previous comparisons (Bradnam et al., 2013). ABySS’ de Bruijn approach did not have issues with the high coverage Illumina data (e.g. Richards & Murali, 2015), which we combined with raw long read data using DGB2OLC (Ye et al. 2016) as previous plant genomes with similar data showed promising assembly statistics (i.e. Hatakeyama et al., 2018, Zhou et al., 2019, Daccord et al., 2017). The long reads in the DBG2OLC assembly captured more lengthy and repetitive regions, resulting in a sharp rise in contig N50 and contigs 50 Kb or longer (Table 1). However, the backbone of the final assembly was really the PacBio Canu assembly (Tables 1 and S4), with the benefit of this strategy being the removal of chimeras with polished assemblies having very low error rates (Ye et al., 2016).

Accurate genomes are important as sequencing errors can affect both nucleotide diversity and the discovery of markers (Clark & Whittam, 1992). Long-read sequencing further facilitates detection of structural variants (Amarasinghe et al., 2020), which might be useful in studying *Typha* dynamics given the link to hybridization and speciation (Weissensteiner et al., 2020). The final assembly consists of only 1,189 scaffolds, with 95.55% being 50 Kb or longer; *T. latifolia* possessed higher N50 and fewer scaffolds compared to other plant genome assemblies that used PacBio sequencing and similar hybrid assembly pipelines (Reuscher et al., 2018, Redwan et al., 2016, Hatakeyama et al., 2018). With 96.03% of BUSCOs detected and 92.28% of them complete in our assembly, this would suggest a high-quality annotation. The topology of the phylogenomic tree that we reconstructed based on whole genome sequences of six species agrees with previously inferred evolutionary relationships (e.g. Darshetkar et al. 2019) by grouping together the two bromeliad species (*T. latifolia* and *A. comosus*) and the three graminid species (*O. sativa, S. bicolor* and *B. distachyon*), and by inferring a more recent divergence date for graminids compared to bromeliads. Our estimated *Typha* lineage age of approximately 70 million years exceeds an earlier estimate of between 22.64 and 57.60 mya that was based on seven cpDNA markers (Zhou et al., 2018), although is comparable to the estimate of 69.5 Myr that was based on a combination of fossil records and cpDNA sequences (Bremer, 2000). We acknowledge that the use of whole genome sequences to infer divergence times is still in its infancy, and caution that factors such as rate heterogeneity may complicate such inferences (reviewed in Smith et al. 2018).

The *T. latifolia* reference genome now allows for studying *Typha* and hybridization at the molecular level, for example by providing insight into the adaptation of *T. latifolia* populations that may be threatened by climate change, and facilitating research into potentially important processes such as hybrid breakdown in invasive *Typha* hybrids. Additionally, genome-wide markers could provide insight into why the hybrid *T*. x *glauca* is dominant in some areas (Kirk et al., 2011; Freeland et al., 2013) but uncommon in others (Freeland et al., 2013; Ciotir et al., 2017). For example, *T. angustifolia* in North America is thought to have arrived from Europe several centuries ago (Ciotir et al., 2013), and ancestry assessments could test the hypothesis that historical interspecific hybridization led to genetic introgression into some *T. angustifolia* populations, which may help to explain regional erosion of species barriers: locus-specific ancestry estimation analyses, markers, and the identification of potentially adaptive genes can all be used to find evidence of ancient hybridization in *T. angustifolia* (Goulet et al., 2017; Taylor & Larson, 2019). This high-quality draft genome and its comparisons with Poales species will be an indispensable resource for ongoing research into *Typha*, a genus that both sustains and threatens wetlands around the world.

## Data Availability Statement

Raw sequencing data can be found in the Short Read Archive (SRA) under accession number PRJNA751759. Genome assembly has been submitted to GenBank (Temporary ID SUB10173221). Code used to generate the data can be found at https://gitlab.com/WiDGeT_TrentU/undergrad-theses/-/tree/master/Widanagama_2021. The *T. angustifolia* RNSseq data is in the SRA under SRR15541138. The *T. latifolia* transcriptome used can be found at https://figshare.com/articles/dataset/Typha_latifolia_leaf_transcriptome/5661727/1.

## Acknowledgements

We thank Vikram Bhargav for sampling and preparing the *T. latifolia* leaves, and Eric Wootton for extracting DNA from the sample. The CanSeq 150 project funded the PacBio sequencing: NSERC Discovery grants to JF and AS and two NSERC Undergraduate Student Research Awards to SW supported this research. XX for the *T. angustifolia* transcriptome used in the genome’s annotation. And Compute Canada Resources for Research Groups award to AS and Compute Canada support staff supported the bioinformatics.

